# Gene neighbourhood integrity disrupted by CTCF loss *in vivo*

**DOI:** 10.1101/187393

**Authors:** Dominic Paul Lee, Wilson Lek Wen Tan, Chukwuemeka George Anene-Nzelu, Peter Yiqing Li, Tuan Anh Luu Danh, Zenia Tiang, Shi Ling Ng, Motakis Efthymios, Matias I. Autio, Jianming Jiang, Melissa Fullwood, Shyam Prabhakar, Roger Sik-Yin Foo

## Abstract

The mammalian genome is coiled, compacted and compartmentalized into complex non-random three-dimensional chromatin loops in the nucleus^1–3^. At the core of chromatin loop formation is CCCTC-binding factor (CTCF), also described as a “weaver of the genome”^45^. Anchored by CTCF, chromatin loops are proposed to form through a loop extrusion process^6^, organising themselves into gene neighbourhoods^2^ that harbour insulated enhancer-promoter domains, restricting enhancer activities to genes within loops, and insulating genes from promiscuous interactions outside of loops^2,7–9^. Studies targeting CTCF binding site deletions at gene neighbourhood boundaries result in localised gene expression dysregulation^8,10–12^, and global CTCF depletion recently showed CTCF to be crucial for higher hierarchical chromatin organisation of topologically associating domains (TADs)^13^. However, the role for CTCF in maintaining sub-TAD CTCF gene neighbourhoods and how gene transcription is affected by CTCF loss remains unclear. In particular, how CTCF gene neighbourhoods govern genome-wide enhancer-promoter interactions require clarification. Here, we took an *in vivo* approach to assess the global dissolution of CTCF anchored structures in mouse cardiomyocyte-specific *Ctcf*-knockout (*Ctcf*-KO), and uncovered large-scale ectopic *de novo* Enhancer-Promoter (E-P) interactions. *In vivo* cardiomyocyte-specific C*tcf*-KO leads to a heart failure phenotype^14^, but our analysis integrates genome-wide transcription dysregulation with aberrant E-P interactions in context of CTCF-loop structures, identifying how genes engage their E-P interactions, requiring CTCF looping for their maintenance. Our study points to a mammalian genome that possesses a strong propensity towards spontaneous E-P interactions *in vivo*, resulting in a diseased transcriptional state, manifest as organ failure. This work solidifies the role of CTCF as the central player for specifying global E-P connections.

## Loss of CTCF causes rapid cardiac dilatation and extensive cardiac gene expression changes

To generate conditional cardiomyocyte-specific *Ctcf*-knockout (*Ctcf*-KO), bypassing early embryonic lethality from *Ctcf*-null^15,16^, and avoiding cardiotoxicity of Tamoxifen in the *Myh6*-MerCremer system^17,18^, we utilized adeno-associated virus 9 (AAV9) to deliver *Cre recombinase*(AAV9:*cTnT*∷*Cre*) specifically targeting cardiomyocytes in the hearts of adult *Ctcf^f/f^* mice^19^. As vector control, we used AAV9:*cTnT*∷*eGFP*(Supplementary Figure 1a, b). Both AAV9-*Cre* and AAV9-*eGFP* cardiac sections showed significant tdTomato and eGFP fluorescence 2 weeks after injection (Supplementary Figure 1c), confirming high transduction efficiency. Cardiac-specific *Ctcf* gDNA deletion was confirmed by PCR analysis (Supplementary Figure 1d). CM-specific CTCF loss was confirmed on histological sections (Supplementary Figure 1e, f), and *Ctcf* mRNA loss was validated by RT-qPCR (Supplementary Figure 1g). Incomplete CTCF depletion may leave residual effects on TAD formation^13,20^, so to further validate the complete absence of CTCF binding on genomic DNA in our model, we performed Western blotting of CM chromatin, validating the complete depletion of chromatin-bound CTCF (Figure 1a). Sustained cardiac dilatation was confirmed starting from 1 week after AAV9:*Cre* injection (Supplementary Figure 1h-j). Dilatation was not accompanied by detectable levels of apoptosis or cell death. Instead, transcriptomic analysis revealed the disarray of important cardiac genes (Supplementary Figure 2a, b; Supplementary Tables 1, 2),including *Ryr2*^21^, *Cacna1c*^22,23^, *Cacnb2*^24^, and *Acta2*^25^, each of which is sufficient as upstream causes to explain cardiac dilatation.

**Figure 1.**
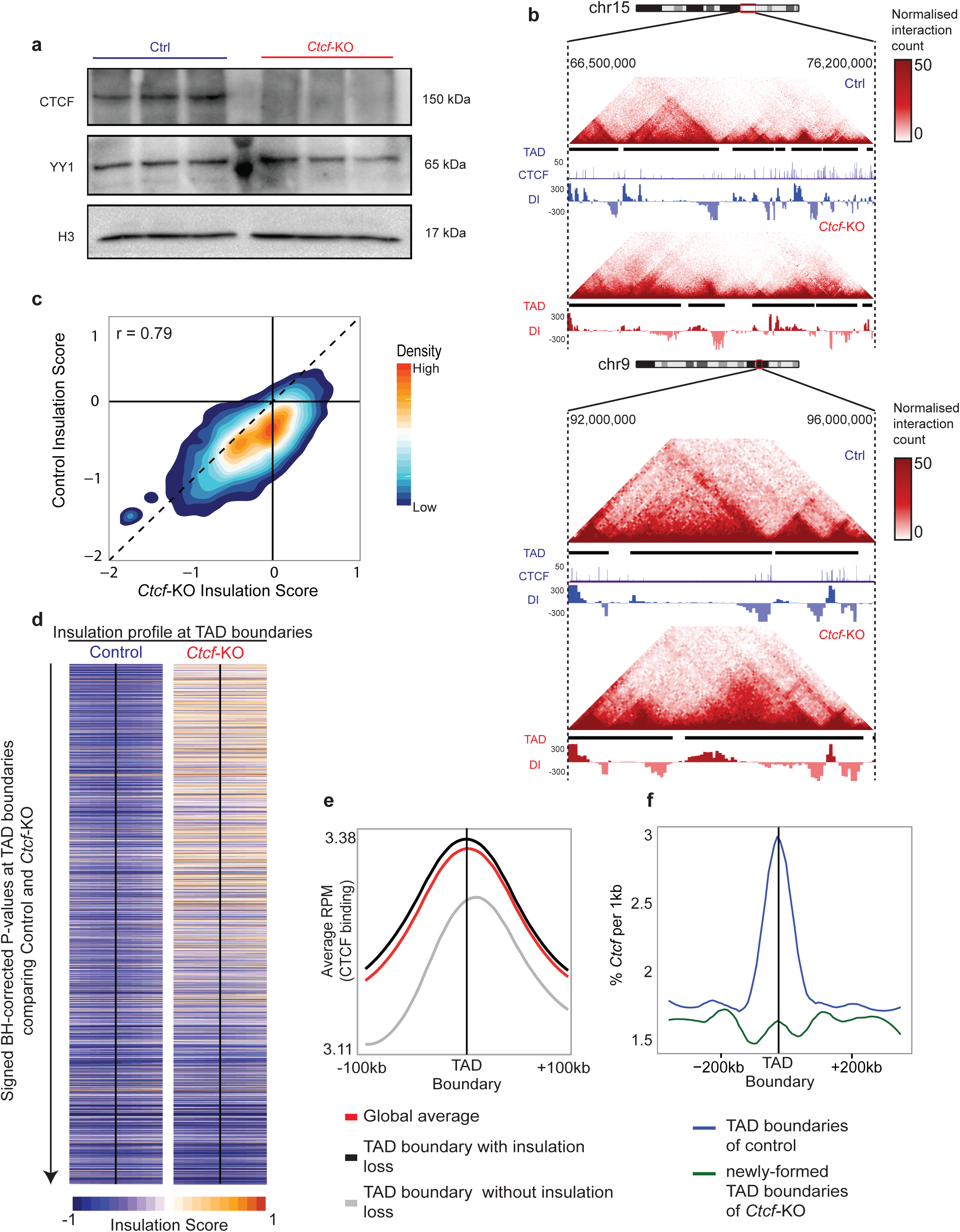
CTCF loss produced severe TAD disruption. **(a)**, Western blot for CTCF protein in cardiomyocyte (CM) chromatin of *Ctcf*-KO and Control. Transcription factor YY1 and histone H3 were used as loading controls. **(b)**, Snapshots of TAD differences between *Ctcf*-KO and Control from Hi-C data identified by Domaincaller^33^. Normalised chromatin interaction heatmaps at 40kb resolution, TADs, and directionality index in *Ctcf*-KO and Control are shown. CTCF ChIP-Seq peaks for Control are also shown. **(c)**, Density plot of genome-wide insulation scores comparing between Control and *Ctcf*-KO. Pearson correlation between insulation scores for *Ctcf*-KO and Control: 0.79; Median score for *Ctcf*-KO and Control: −0.27, −0.54, respectively. Wilcoxon rank sum test with continuity correction, *P*<2.2x10^−16^) (**d)**, Insulation scores at genome-wide TAD boundaries (flanking 100kb) ranked according to median insulation score difference between Control and *Ctcf*-KO. **(e)**, CTCF occupancy at global TAD boundaries with and without loss of insulation scores, based on CTCF ChIP-seq in Control CMs. Significantly higher CTCF occupancy was found for TAD boundaries with insulation loss (N=2,089), compared to those without insulation loss (N=439) (*P*< 2.2x10^−16^; Wilcoxon rank sum test with continuity correction). **(f)**, CTCF occupancy at genome-wide TAD boundaries of Control and newly-formed TAD boundaries in *Ctcf*-KO, based on CTCF ChIP-seq in Control.

Progressive heart failure is itself characterised by a stress-gene response, typified by the re-expression of fetal cardiac genes^26,27,28^. While it may be possible that *Ctcf*-KO expression changes are the result of progressive disease rather than a direct effect of CTCF loss itself, we noted that *Ctcf*-KO hearts in fact recapitulated little of the classical disease stress-gene response (Supplementary Figure 2c)^26^-^28^, suggesting that at least a part of the expression change arises from CTCF perturbation, and not simply from disease progression.

## Loss of CTCF disrupts TADs and sub-TAD gene neighbourhoods

To investigate for 3D chromatin organisation genome-wide, we performed *in-situ* Hi-C^1^ on ventricular CMs that were isolated using our published protocol^29^ from *Ctcf*-KO and Control mice. Hi-C libraries were sequenced to a depth of 575M and 589M read pairs, respectively. Replicate libraries were combined to attain deeper coverage (Supplementary Table 1).

First, we analysed for megabase-scale TADs^30,31^ by using Domain-Caller to identify TAD boundaries systematically based on directionality score. 2,199 and 1,777 TADs were identified in Control and *Ctcf*-KO (Figure 1b, Supplementary Figure 3, Supplementary Table 3). TAD number decrease was also accompanied by an increase in genome-wide TAD lengths, likely reflecting a weakening or dissolution of TAD boundaries, and consequent TAD widening (1.0 ± 0.81Mb, 1.28 ± 0.84Mb for Control and *Ctcf*-KO TAD lengths respectively; *p*<2.2x10^−16^; Wilcoxon sum rank test). To confirm the phenomenon of weakened TAD boundaries, we computed for genome-wide insulation scores^32^, where higher scores represent loss of insulation integrity. *Ctcf*-KO led to higher insulation scores (Figure 1c,d), reflecting the loss of insulation at TAD boundaries and their overall dependence on CTCF, as also reported recently^13^. Since we had assessed CTCF to be completely depleted in our CM chromatin (Figure 1a), this also gave us the opportunity to further analyse what, apart from CTCF, characterised remaining TAD boundaries in *Ctcf*-KO. For reference, we generated triplicate CTCF ChIP-seq using isolated CMs from Control (Supplementary Table 4). From these, we saw, as also previously reported^33^, that CTCF was enriched at only ~75-80% of Control TAD boundaries. Indeed, in *Ctcf*-KO, TAD boundaries with insulation loss were significantly enriched for CTCF occupancy, compared to boundaries without insulation loss (*P*<2.2x10^−16^; (Figure 1e). In fact, TAD boundaries in Control were CTCF-enriched, whereas newly-acquired boundaries in *Ctcf*-KO were not (Figure 1f). Moreover, boundaries in *Ctcf*-KO lacked other characteristic features of TAD boundaries compared to Control (Supplementary Figure 4). Apart from an enrichment of gene transcription start sites (TSS) and adjacent RNApol2 binding sites, this analysis gave an overall unclear impression of where boundaries dissolved to following CTCF loss.

As only 20% of genome-wide CTCF binding sites localised to TAD boundaries^1,31^(Supplementary Figure 5), and the majority of CTCF are instead localised to within TADs, we turned our next analysis towards CTCF-anchored loops including those inside TADs. To this end, we partitioned the mouse genome into 10kb bins, and curated for statistically significant *bona fide* genome-wide chromatin interactions using Fit-Hi-C (Figure 2a, b)^34,35^ (Supplementary Table 5). Most bins (85%) annotated with TSS were occupied by one unique TSS; and similarly, most H3K27ac-annotated (70%) or CTCF-annotated (80%) bins contained at most one H3K27ac peak or one CTCF peak respectively, allowing us to use our dataset to study CTCF-CTCF and Enhancer-Promoter (E-P) interactions at high resolution. Overall, intra-chromosomal contacts were independently identified in Control (125,079) and *Ctcf*-KO (97,789) (FDR<0.01, normalized contact>10) (Figure 2b). Of those in Control, only 25,664 remained in *Ctcf*-KO, and 72,125 were new. Overall, 18,725 were CTCF-CTCF interactions in Control, reduced to 5,521 in *Ctcf*-KO, reflecting an overall CTCF-CTCF loss following CTCF deletion (Supplementary Table 5). Of the 18,725 interactions in Control, we noticed that some CTCF anchors also contained Enhancers and Promoters (Supplementary Figure 6). Hence, to define CTCF-CTCF loop loss more precisely, we prioritised the anchors for Promoter > Enhancer > CTCF, as also performed in Ji *et al*^10^. Applying this strategy, we found 7,224 CTCF-CTCF interactions where both anchors contained only CTCF in Control, and this number was again reduced to 1,572 in *Ctcf*-KO (Figure 2c). Even so, to further cross-validate for CTCF-loop loss, we employed an independent loop-calling tool, HICCUPS^1,36^, and observed a 97% global loss of CTCF-CTCF contacts (2,908 and 103 loops in Control and *Ctcf*-KO, respectively, FDR<0.1; Figure 2d; Supplementary Table 6). Of note, almost all loops called by HICCUPS were also identified by Fit-Hi-C (Supplementary Figure 7). Taken together, global CTCF-CTCF looping interactions were indeed largely abrogated upon CTCF deletion in our model.

**Figure 2.**
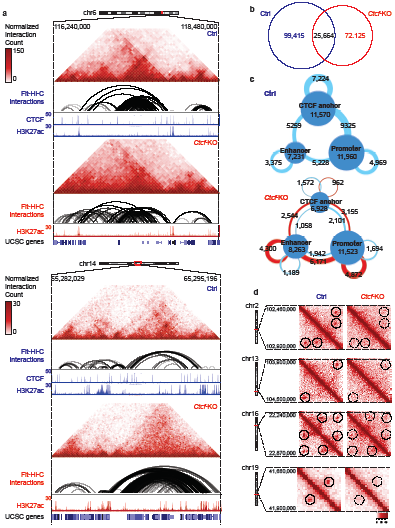
CTCF loss produced significant loss of Hi-C interactions, as well as widespread gain of ectopic interactions. **(a)**, Snapshots of Hi-C interaction data at 10kb resolution in Control and *Ctcf*-KO. Normalised Hi-C heatmaps, significant Fit-Hi-C interactions (FDR<0.01), H3K27ac and CTCF ChIP-seq for loci on chromosomes 6 and 14 are shown. **(b)**, Venn diagram of high confidence interaction counts (FDR < 0.01) from *Ctcf*-KO and Control Hi-C. **(c)**, Significant losses and gains in interactions between CTCF, Enhancers (H3K27ac) and Promoter (TSS) anchors comparing between Control and *Ctcf*-KO, based on CTCF ChIP-seq in Control and H3K27ac ChIP-seq in Control and *Ctcf*-KO. Circles represent anchors based on 10kb windows, assigned as either promoter, enhancer or CTCF, based on the order of priority (Promoter > Enhancer > CTCF). Connecting ribbons depict interactions between anchors. Size of circles is proportional to the number of anchors, and thickness of ribbons is proportional to the number of interactions. Red ribbons represent *de novo* interactions gained in *Ctcf*-KO. **(d)**, Near complete dissolution of loops (circled) in *Ctcf*-KO, as identified by HICCUPS analysis (FDR<0.1).

## Widespread promiscuous E-P interactions in the Ctcf-KO genome

Previous studies using Crispr-deletion of specific CTCF binding sites resulted in localised gene dysregulation near deleted loop boundaries, underpinning the structure of CTCF-insulated gene neighbourhoods^8,10,11,37–39^. We therefore asked if chromatin loop loss causes gene dysregulation by *de novo* Enhancer-Promoter (E-P) interactions, and if so, whether they extended across gene neighbourhoods. High-confidence interactions were retained for only 1,942 of 5,228 E-P interactions comparing between Control and *Ctcf*-KO. Strikingly, this accompanied 6,171 new *de novo* E-P interactions in *Ctcf*-KO, and similar increases in enhancer-enhancer and promoter-promoter interactions (Figure 2c).

Next, we computed directionality index at 10kb resolution and annotated 2,457 insulated CTCF-CTCF loops (Figure 3a-c, Table and Supplementary Table 7). In order to prove the validity of our insulated CTCF neighbourhoods, we confirmed that E-P interactions starting with an anchor in a neighbourhood predominantly ended within the confines of the neighbourhood, supporting the conclusion that CTCF-loops form insulated gene neighbourhoods and prevent cross-boundary interactions (Figure 3a, Supplementary Figure 8)^8,10,11,40^. In contrast, new *de novo* E-P interactions at the corresponding loci in *Ctcf*-KO showed clear evidence of boundary crossing (Figure 3b, c). Overall, only 33% (394/1,190) of E-P interactions were cross-boundary in Control, whereas this was increased to 73% (1,011/1,384) in *Ctcf*-KO (*P*<2.2x10^−16^, Fisher’Mxtables exact test). Moreover, the majority of new *de novo* E-P interactions in *Ctcf*-KO utilised pre-existing active enhancers in Control, now acting with new promoter sites (Figure 3d). Coherent with the notion that lost loop boundaries result in *de novo* E-P interactions that induce gene upregulation, and loss of intra-loop E-P interactions induces gene down-regulation, we found that indeed, upregulated genes were enriched both inside loops and extended outwards of lost loop boundaries, whereas down-regulated genes were mainly confined to inside loops (Figure 3e). Moreover, gained E-P interactions correlated globally with gene up regulation, and vice versa (Figure 3f).

**Figure 3.**
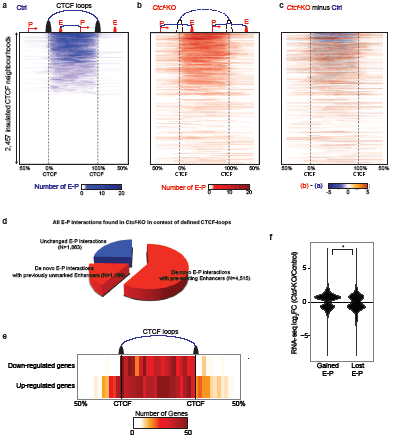
CTCF loss disrupted insulated gene neighbourhoods, producing global cross boundary E-P interactions, and consequent gene expression changes. **(a)**, E-P interactions are largely confined within genome-wide CTCF-bound insulated gene neighbourhoods in Control. Each row represents an insulated CTCF neighbourhood. Columns represent scaled distance (0-100%) from the left CTCF anchor, for both inside and outside of the CTCF-CTCF neighbourhood. Intensity of blue horizontal lines represents number of E-P interactions covering the locus, where the E-P interaction has at least one anchor inside the CTCF-CTCF neighbourhood. **(b)**, E-P interactions at the same loci in (a) now in *Ctcf*-KO, many of which span across previously confined CTCF boundaries. As in (a), intensity of green horizontal lines represent E-P interactions with at least one anchor inside the previously defined CTCF-CTCF neighbourhood. (**c**), The subtraction of E-P interactions between (b) *Ctcf*-KO and (a) Control. Newly-formed E-P interactions in *Ctcf*-KO are depicted in orange, and lost E-P interactions in blue. **(d)**, Breakdown of E-P interactions found in *Ctcf*-KO. 1,863 were E-Ps also found in Control. The rest were new *de novo* E-Ps, of which 4,515 were ones that utilised pre-existing active Enhancers also seen in Control. **(e)**, Location of differently expressed genes in context of CTCF-CTCF neighbourhoods. Up-regulated genes (fold-change >1.3, >4 FPKM) in *Ctcf*-KO mice were found both inside and outside of CTCF-CTCF neighbourhoods, whereas down-regulated genes were mainly confined inside CTCF neighbourhoods (Fisher’s exact test *P*=9.3x10^−7^, odds ratio of observing upregulated genes outside CTCF neighbourhood =1.75). **(f)**, Gained E-P interactions in *Ctcf*-KO correlated with up-regulated gene expression, and vice versa for lost E-P interactions (Wilcoxon rank sum test with continuity correction *P*=0.01).

**Table.** Characteristics of CTCF gene neighbourhoods (N=2,457)

**Table. Characteristics of CTCF gene neighbourhoods (N=2,457)**
| Lengths of CTCF gene neighbourhoods |  |  |  |
| --- | --- | --- | --- |
| Min: 60kb | Median: 370kb | Mean: 462kb | Max: 3.3Mb |
| Gene distribution inside CTCF neighbourhoods |  |  |  |
| Min: 0 gene | Median: 5 genes | Mean: 7 genes | Max: 103 genes |
| Enhancer distribution inside CTCF neighbourhoods |  |  |  |
| Min: 0 | Median: 6 | Mean: 9 | Max: 89 |

## A cardiac gene neighbourhood exemplifies context-dependent gene dysregulation and interaction gain and loss

Specifically to cardiac stress-response genes, we noted the divergent expression change of 2 important disease markers - *Nppa* and *Nppb*^28,29,41^ (Figure 4a). In *Ctcf*-KO, *Nppb* was upregulated, whereas *Nppa* was not, raising the possibility that the difference may relate to a disrupted gene neighbourhood. To confirm that this was a direct consequence of *Ctcf*-KO and not due to the lack of a sufficient stress response, we further induced surgical pressure-overload by transverse aortic constriction (TAC) on *Ctcf*-KO mice, and confirmed the lack of stress-response in *Nppa*, but not *Nppb*(Figure 4a).

**Figure 4.**
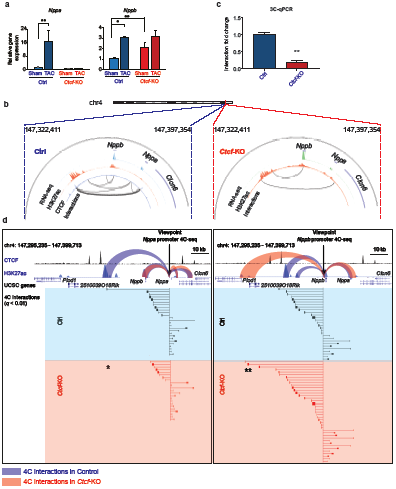
An example of chromatin interactions at the *Nppa*/*Nppb* stress-gene locus. **(a)**, RT-qPCR results of *Nppa* and *Nppb* stress-response in Control and *Ctcf*-KO, without and with cardiac pressure-overload (sham and TAC, respectively). Data represent mean ± s.e.m. \**P*<0.05, \*\**P*<0.01. Student’s t test. **(b)**, Circos plots for Hi-C interactions within the CTCF-CTCF loop containing *Nppa* and *Nppb.***(c)**, 3C-qPCR for the CTCF-CTCF loop interaction depicted in (b). Data represents mean ± s.e.m. \*\**P*<0.01. Student’s t test. **(d, e)**, 4C-seq for the *Nppa* (d), and *Nppb* promoters (e), showing the loss of interaction to an upstream element (*) for *Nppa* in *Ctcf-KO*,but not for *Nppb*. Instead, *Nppb* acquires a *de novo* enhancer-interaction (**) further upstream and outside of the CTCF anchor. CTCF ChIP-seq was performed for Control only, whereas H3K27ac ChIP-seq was for both and *Ctcf*-KO.

*Nppa* and *Nppb* both reside in a CTCF-CTCF anchored gene neighbourhood, flanked by convergent CTCF sites (Figure 4b)^42^. We validated the loss of this gene neighbourhood in *Ctcf-*KO, by an 8-fold decrease in the CTCF-CTCF interaction located 49kb apart using 3C-qPCR (Figure 4c). Next, we performed 4C-seq using the *Nppa* and *Nppb* promoters separately as viewpoints (Figure 4d, Supplementary Tables 8, 9). All significant interactions involving *Nppa* and *Nppb* were indeed confined to within the CTCF-anchored neighbourhood. Both genes interacted with an upstream H3K27ac-enriched region, 34kb from *Nppa*(vertical red box in (Figure 4d), also previously implicated as a putative *Nppa*/*Nppb* enhancer^28,42^. In *Ctcf*-KO, this upstream E-P interaction with *Nppa* alone was abolished. The interaction loss was associated with the lack of *Nppa* stress-gene response, compared to the intact *Nppb* response. Remarkably, accompanying the loss of this insulated neighbourhood in *Ctcf*-KO, we also serendipitously noted the gain of a promiscuous *de novo* interaction for the *Nppb* promoter with a further upstream gene *Plod1*(48kb away), outside of the previously demarcated CTCF boundary (black box in (Figure 4d, Supplementary Figure 9). Furthermore, this *de novo* interaction was also accompanied by significant upregulation of *Plod1* expression (Supplementary Figures 2, 10).

## CONCLUSION

Our study utilising Hi-C datasets after CTCF deletion in CMs underscores CTCF as central to the formation of gene neighbourhoods and provides evidence for CTCF loops as basal units of genomic loop organization. Insulated neighbourhoods not only confine E-P interactions to within themselves, but are also crucial for facilitating intra-loop interactions, as in the case of *Nppa* gene. Conversely, the mammalian genome is susceptible to the formation of ectopic enhancer interactions that in turn misregulate gene expression when not constrained in gene neighbourhoods as dictated by CTCF. This work brings into question whether true specificity of enhancers with their target promoters exists, and instead describes a genome where transcription is the consequence of E-P interactions dependent on their encasement in CTCF loops. Future studies on cell type specific gene neighbourhoods and active enhancers within, will be needed to uncover genetic variants that influence gene looping and human diseases.

## Author contributions

D.P.L., C.G.A-N., W.L.W.T., S.L.N., Z.T., J.J. and M.I.A. performed the experiments, collected the data and carried out the analysis. P.Y.Q. and T.A.L.D. carried out the animal procedures. M.E. and W.L.W.T. performed the bioinformatics and computational analyses. M.E. performed GO analysis. M.I.A., M.F., S.P. and R.S-Y.F supervised the project. D.P.L., C.G.A-N., W.L.W.T., and R.S-Y.F. wrote the manuscript.

## Acknowledgement

This project was funded by grants from the Biomedical Research Council (BMRC), Agency for Science, Technology and Research (A*STAR) and National Medical Research Council (NMRC) of Singapore. R.S-Y.F. was funded by a CSA (SI) award. D.P.L. was funded by an NUS Research Scholarship. We thank Dr Ong Chin Tong (Temasek Life Sciences Lab, Singapore) for critical reading of the manuscript.

## References

1. Rao, S. S. P. et al. A 3D map of the human genome at kilobase resolution reveals principles of chromatin looping. Cell 159, 1665–1680 (2014).

2. Hnisz, D., Day, D. S. & Young, R. A. Insulated Neighborhoods: Structural and Functional Units of Mammalian Gene Control. Cell 167, 1188–1200 (2016).

3. Bonev Boyan & Cavalli Giacomo. Organization and function of the 3D genome. Nat. Rev. Genet. 17, 661–678 (2016).

4. Ong, C.T. & Corces, V. G. CTCF: an architectural protein bridging genome topology and function. Nat. Rev. Genet. 15, 234–46 (2014).

5. Phillips, J. E. & Corces, V. G. CTCF: master weaver of the genome. Cell 137, 1194–211 (2009).

6. Fudenberg, G. et al. Formation of Chromosomal Domains by Loop Extrusion. Cell Rep. 15, 2038–2049 (2016).

7. Tang, Z. et al. CTCF-Mediated Human 3D Genome Architecture Reveals Chromatin Topology for Transcription. Cell 163, 1611–1627 (2015).

8. Dowen, J. M. M. et al. Control of Cell Identity Genes Occurs in Insulated Neighborhoods in Mammalian Chromosomes. Cell 159, 374–387 (2014).

9. Phillips-Cremins, J. E. et al. Architectural protein subclasses shape 3D organization of genomes during lineage commitment. Cell 153, 1281–95 (2013).

10. Ji, X. et al. 3D Chromosome Regulatory Landscape of Human Pluripotent Cells. Cell Stem Cell 18, 262–275 (2016).

11. Hnisz, D. et al. Activation of proto-oncogenes by disruption of chromosome neighborhoods. Science 351, 1454–1458 (2016).

12. Flavahan, W. A. et al. Insulator dysfunction and oncogene activation in IDH mutant gliomas. Nature 529, 110–114 (2015).

13. Nora, E. P. et al. Targeted Degradation of CTCF Decouples Local Insulation of Chromosome Domains from Genomic Compartmentalization. Cell 169, 930–944.e22 (2017).

14. Rosa-Garrido, M. et al. High Resolution Mapping of Chromatin Conformation in Cardiac Myocytes Reveals Structural Remodeling of the Epigenome in Heart Failure. Circulation CIRCULATIONAHA.117.029430 (2017). doi:10.1161/CIRCULATIONAHA.117.029430

15. Heath, H. et al. CTCF regulates cell cycle progression of alphabeta T cells in the thymus. EMBO J. 27, 2839–50 (2008).

16. Fedoriw, A. M., Stein, P., Svoboda, P., Schultz, R. M. & Bartolomei, M. S. Transgenic RNAi reveals essential function for CTCF in H19 gene imprinting.Science 303, 238–40 (2004).

17. Bersell, K. et al. Moderate and high amounts of tamoxifen in αMHC-MerCreMer mice induce a DNA damage response, leading to heart failure and death. Dis. Model. Mech. 6, 1459–69 (2013).

18. Werfel, S. et al. Rapid and highly efficient inducible cardiac gene knockout in adult mice using AAV-mediated expression of Cre recombinase. Cardiovasc. Res. 15–23 (2014). doi:10.1093/cvr/cvu174

19. Lin, Z. et al. Cardiac-Specific YAP Activation Improves Cardiac Function and Survival in an Experimental Murine MI Model. Circ. Res. 115, 354–363 (2014).

20. Zuin, J. et al. Cohesin and CTCF differentially affect chromatin architecture and gene expression in human cells. Proc. Natl. Acad. Sci. U. S. A. 111, 996–1001 (2014).

21. Bround, M. J. et al. Cardiac ryanodine receptors control heart rate and rhythmicity in adult mice. Cardiovasc. Res. 96, 372–380 (2012).

22. Goonasekera, S. A. et al. Decreased cardiac L-type Ca 2 + channel activity induces hypertrophy and heart failure in mice. J. Clin. Investig.http://www.jci.org, 122 (2012).

23. Rosati, B. et al. Robust L-type calcium current expression following heterozygous knockout of the Cav1. 2 gene in adult mouse heart. J. Physiol. 589, 3275–88 (2011).

24. Meissner, M. et al. Moderate calcium channel dysfunction in adult mice with inducible cardiomyocyte-specific excision of the cacnb2 gene. J. Biol. Chem. 286, 15875–15882 (2011).

25. Schildmeyer, L. a et al. Impaired vascular contractility and blood pressure homeostasis in the smooth muscle alpha-actin null mouse. FASEB J. 14, 2213–2220 (2000).

26. See, K. et al. Single cardiomyocyte nuclear transcriptomes reveal a lincRNA-regulated de-differentiation and cell cycle stress-response in vivo. Nat. Commun. 8, 225 (2017).

27. Lee, J.-H. et al. Analysis of Transcriptome Complexity Through RNA Sequencing in Normal and Failing Murine Hearts. Circ. Res. 109, 1332–1341 (2011).

28. Dirkx, E., da Costa Martins, P. a. & De Windt, L. J. Regulation of fetal gene expression in heart failure. Biochim. Biophys. Acta - Mol. Basis Dis. 1832,2414–2424 (2013).

29. Ackers-Johnson, M. et al. A Simplified, Langendorff-Free Method for Concomitant Isolation of Viable Cardiac Myocytes and Nonmyocytes from the Adult Mouse Heart. Circ. Res. 119, 909–920 (2016).

30. Nora, E. P. et al. Spatial partitioning of the regulatory landscape of the X-inactivation centre. Nature 485, 381–385 (2012).

31. Dixon, J. R. et al. Topological domains in mammalian genomes identified by analysis of chromatin interactions. Nature 485, 376–380 (2012).

32. Crane, E. et al. Condensin-driven remodelling of X chromosome topology during dosage compensation. Nature 523, 240–244 (2015).

33. Dixon, J. R. et al. Topological domains in mammalian genomes identified by analysis of chromatin interactions. Nature 485, 376–80 (2012).

34. Ay, F., Bailey, T. L. & Noble, W. S. Statistical confidence estimation for Hi-C data reveals regulatory chromatin contacts. Genome Res. 24, 999–1011 (2014).

35. Wang, M. et al. Asymmetric subgenome selection and cis-regulatory divergence during cotton domestication. Nat. Genet. 49, 579–587 (2017).

36. Durand, N. C. et al. Juicer Provides a One-Click System for Analyzing Loop-Resolution Hi-C Experiments. Cell Syst. 3, 95–98 (2016).

37. Hanssen, L. L. P. et al. Tissue-specific CTCF–cohesin-mediated chromatin architecture delimits enhancer interactions and function in vivo. Nat. Cell Biol. 19, (2017).

38. Narendra, V. et al. CTCF establishes discrete functional chromatin domains at the Hox clusters during differentiation. Science 347, 1017–1021 (2015).

39. Lupiáñez, D. G. et al. Disruptions of topological chromatin domains cause pathogenic rewiring of gene-enhancer interactions. Cell 161, 1012–1025 (2015).

40. Phillips-Cremins, J. E. & Corces, V. G. Chromatin Insulators: Linking Genome Organization to Cellular Function. Mol. Cell 50, 461–474 (2013).

41. Creemers, E. E., Wilde, A. a & Pinto, Y. M. Heart failure: advances through genomics. Nat. Rev. Genet. 12, 357–62 (2011).

42. Sergeeva, I. A. et al. Identification of a regulatory domain controlling the Nppa-Nppb gene cluster during heart development and stress. Development 143, 2135–2146 (2016).

